# Bicistronic expression of a high-performance calcium indicator and opsin yields stable, robust cortical expression for holographic two-photon stimulation

**DOI:** 10.1101/2022.09.06.506779

**Authors:** Paul K. LaFosse, Zhishang Zhou, Nina G. Friedman, Yanting Deng, Bradley Akitake, Mark H. Histed

## Abstract

Patterns of activity across many neurons are fundamental units of neural computation. Two-photon holographic photostimulation allows both delivering input to, and imaging responses from, patterns or populations of neurons. However, to make this method an easily-deployable tool, simple methods are needed to robustly and stably express opsins and indicators in the same cells. Here we describe a bicistronic adeno-associated virus (AAV) that in transfected cells expresses both the fast and bright calcium indicator GCaMP8s, and a soma-targeted (st) and two-photon-activatable opsin, ChrimsonR. With this method, in the visual cortex of mice, stChrimsonR stimulation with two-photon holography drives robust spiking in targeted cells, and neural responses to visual sensory stimuli and spontaneous activity are strong and easy to measure. stChrimsonR is a good choice of opsin when a balance is needed between stimulation-laser activatability and avoidance of imaging laser activation. This approach is a simple and robust way to prepare neurons *in vivo* for two-photon holography and imaging.

**Significance statement:** The recent advent of holographic photostimulation methods in conjunction with standard two-photon calcium imaging promises unprecedented levels of control in manipulating and dissecting brain circuitry *in vivo* while reading out neural activity. These all-optical methods rely on a working synergy between optogenetic strategies to both measure calcium activity through genetically-encoded calcium indicators and modulate cell activity through light-activated opsins. Genetic strategies to achieve reliable and stable co-expression of opsin and indicator remain sparse and often challenging to execute. Here, we present a genetic tool to achieve robust co-expression of jGCaMP8s indicator and stChrimsonR opsin via a single injected virus to help facilitate experiments aiming to use holography to investigate the circuit principles underlying brain activity.

## Introduction

Perception and action depend on neural computations, created as activity patterns propagate in neural circuits. Monitoring and controlling these activity patterns are an essential step in understanding how brain function governs perception and behavior. Optogenetics has made it widely possible to control genetically-specified sets of neurons. However, achieving optical specificity — the ability to select a single cell and perturb it — is challenging. Conventional one-photon excitation methods are not ideal, as light can penetrate hundreds of microns in depth throughout the tissue, resulting in undesired activation outside of the focal plane. Moreover, one-photon methods are limited in their ability to restrict excitation to small volumes due to light scattering in the tissue (Denk et al., 1994).

Two-photon optogenetics overcomes these limitations to enable perturbations in selected single cells (Rickgauer and Tank, 2009; Packer et al., 2012, 2015; Emiliani et al., 2015; Adesnik and Abdeladim, 2021). With this approach, using a stimulation and an imaging laser independently focused at different locations deep in the brain, it is possible to simultaneously measure evoked activity patterns as stimulation is delivered. Two-photon optogenetics has been used to study within-area network dynamics (Chettih and Harvey, 2019), and to understand how chosen patterns of activity evoked by stimulation influence perception (Carrillo-Reid et al., 2019; Marshel et al., 2019; Dalgleish et al., 2020; Gill et al., 2020; Robinson et al., 2020; Daie et al., 2021; Rowland et al., 2021; Russell et al., 2022).

A variety of opsins and calcium indicators have been used for two-photon stimulation and simultaneous imaging (Shemesh et al., 2017; Mardinly et al., 2018; Chen et al., 2019; Marshel et al., 2019; Adesnik and Abdeladim, 2021; Forli et al., 2021; Sridharan et al., 2022). Desirable properties for calcium indicators used with two-photon stimulation include high sensitivity to measure small changes in neural firing, and fast dynamics to monitor fast-changing spike trains. Desirable properties for opsins include fast dynamics to allow precise control of spiking and activation using moderate to low stimulation intensities, which allows many neurons to be stimulated without excessive energy being applied to the brain.

It has proven challenging to co-express opsins and calcium indicators in the same neurons stably for weeks to months. High levels of indicator expression in single cells can lead to reduced fluorescence responses, typically with constant levels of bright fluorescence, a phenomenon that can become more common as time elapses after transfection (Tian et al., 2009; Chen et al., 2013; Packer et al. 2015). Further, when expressing both proteins via separate viruses, it can be challenging to achieve co-expression in many neurons (see Packer et al., 2015; Carrillo-Reid et al. 2018, their Fig. 4a; Chettih and Harvey, 2019; Gill et al., 2020; Russell et al., 2022). Genetic mouse lines promise to simplify this co-expression process (Bounds et al., 2022), but current genetic lines have limited combinations of opsin and calcium indicator available, and in genetic lines it has not always been possible to achieve the levels of indicator expression (Daigle et al., 2018) that give imaging quality comparable to viral expression.

Here we demonstrate a single Cre-dependent virus that expresses both opsin and indicator in transfected cells without requiring multiple overlapping viral injections. Our solution uses the ChrimsonR opsin, targeted to cells’ somatas with a Kv2.1 domain (soma-targeted ChrimsonR, stChrimsonR, Pégard et al., 2017), and jGCaMP8s, a bright, sensitive genetically encoded calcium indicator (Zhang et al., 2021). The genes are linked by the self-cleaving peptide P2A (Szymczak et al., 2004; Prakash et al., 2012), an approach previously used with GCaMP6m and the opsin ChRmine (Marshel et al. 2019). stChrimsonR is an opsin with fast on- and off-kinetics (Klapoetke et al., 2014) with a red-shifted excitation spectrum (see two-photon excitation spectrum in Supp Fig. 3 of Sridharan et al., 2022) and moderate sensitivity to two-photon activation (Chen et al., 2019). This moderate sensitivity to stimulation has the advantage of allowing neurons to be driven by the stimulation laser, but avoiding activation, or crosstalk, from the (lower peak intensity) imaging laser. jGCaMP8s is from the latest generation of fast calcium indicators, balancing needs for a bright signal and physiologically relevant kinetics. We find this construct provides a stable preparation for long-term experiments with repeated stimulation. With this single virus strategy, we achieve widespread and stable expression, effective and precise holographic stimulation of many cells, and high-quality recording of neural activity.

## Results

We made a single adeno-associated virus (AAV; Fig. 1A) that contains the genes for jGCaMP8s (Zhang et al., 2021) and ChrimsonR (Klapoetke et al., 2014), separated by the P2A cleavage site, which allows for similar ratios of expression of both proteins (Kim et al., 2011) from cell to cell. The Cre dependence is provided by the DIO (FLEX) strategy (Schnütgen et al., 2003; Cardin et al., 2009), such that without recombination, the genes are in the antisense orientation to limit leaky expression in the absence of Cre.

**Figure 1:**
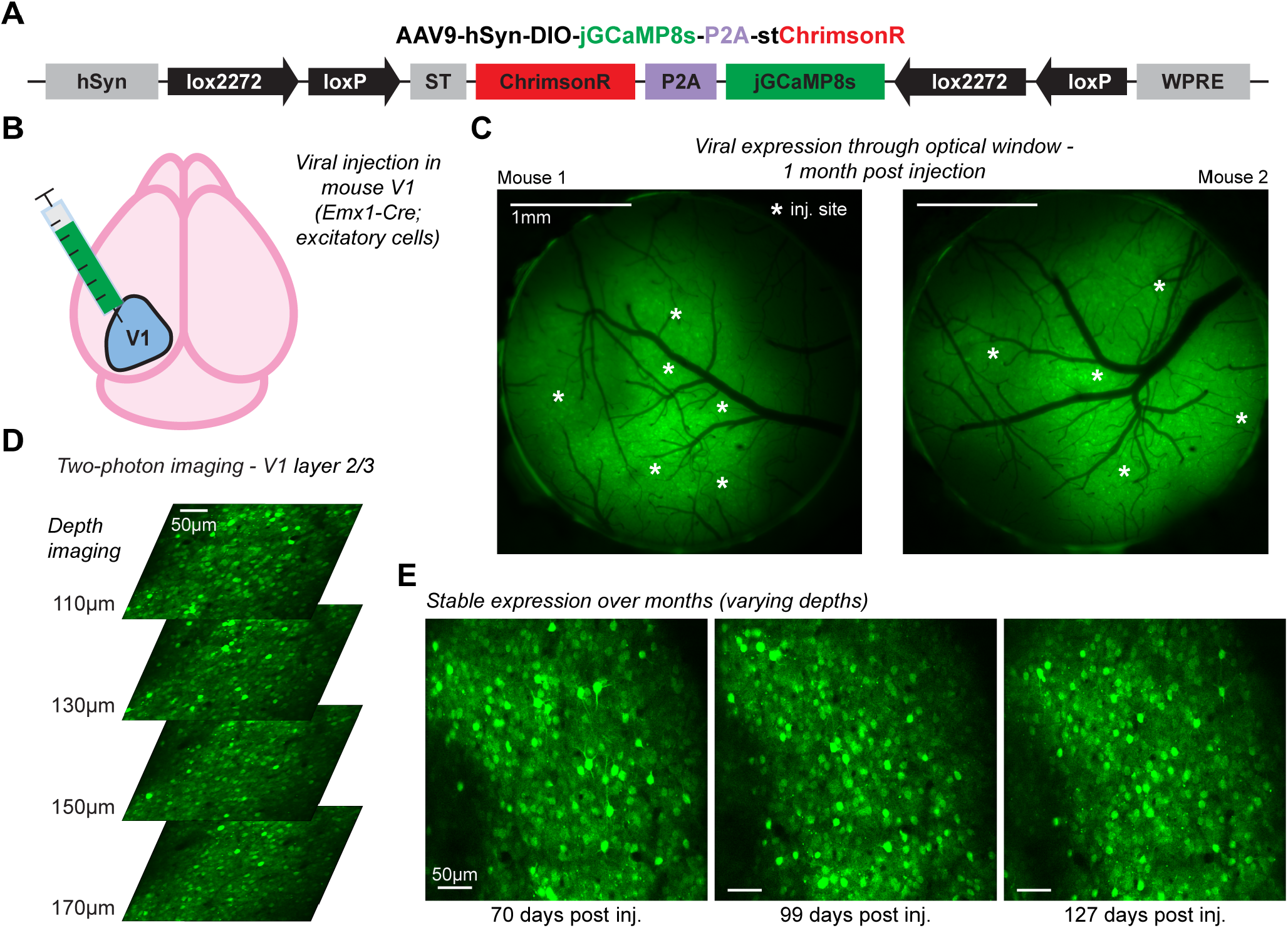
Widespread and stable co-expression of jGCaMP8s and stChrimsonR in mouse cortical neurons with a bicistronic viral vector. ***A*** Schematic of the Cre-dependent, bicistronic vector. ***B*** Viral injections were administered to primary visual cortex (V1) to achieve targeted expression in excitatory populations using an *Emx1-Cre* mouse line. ***C*** Widefield fluorescence imaging through 3 mm optical windows in two example animals shows widespread expression over the cortical surface surrounding injection sites. ***D*** Two-photon calcium imaging (at 920 nm) in mouse V1 throughout layer 2/3 shows robust expression of the virus across many cells (FOV 410 × 410 μm). ***E*** Repeated two-photon imaging in the same area reveals stable expression over many months by returning to approximately the same FOV multiple times in one animal (with numerous two-photon imaging and optogenetic stimulation experiments occurring between; N = 5 stimulation experiments, N = 6 total imaging experiments). This figure: N = 3 mice. **C**: mouse 1-2, **D**-**E**: mouse 3.

### Robust and stable expression of jGCaMP8s and stChrimsonR using a bicistronic construct

For all experiments, we injected the virus into primary visual cortex (V1) of adult *Emx1-Cre* mice, to yield expression in excitatory neurons (Fig. 1B). We implanted optical windows over V1 for imaging.

At three weeks post-injection, strong GCaMP fluorescence was visible using fluorescence imaging of the cortical surface through the optical window (N = 6 animals injected and measured 22-53 days post-injection; Fig. 1C and Extended Fig. 1-1). To assess expression in individual cells, we used *in vivo* two-photon calcium imaging. We found robust expression in many cells across multiple imaging depths (Fig. 1D). We also found stable expression over several months (Fig. 1E), even though multiple stimulation experimental sessions were interspersed between the imaging sessions.

We find that many or all neurons show GCaMP fluorescence throughout the cell, unlike what is seen with expression of GCaMP alone under control of a single promoter, where GCaMP is often excluded from the nucleus (Tian et al., 2009; Chen et al., 2013; Packer et al. 2015). While this might imply some differences in GCaMP trafficking with this bicistronic virus compared to GCaMP expressed with a single promoter, we next characterize responses to both visual stimulation and holographic stimulation and find robust responses in both cases.

### In vivo recording of visually-evoked and spontaneous activity of V1 cells expressing jGCaMP8s-P2A-stChrimsonR

In order to determine whether cells expressing jGCaMP8s-P2A-stChrimsonR exhibit physiologically expected sensory-evoked responses (see Kondo and Ohki, 2016; Zhang et al., 2021; Bounds et al., 2022), we used two-photon imaging to record jGCaMP8s activity from excitatory cells in layer 2/3 of mouse V1 during and between presentations of retinotopically-aligned drifting grating (Gabor patches, 15 degree full-width half-max, FWHM) stimuli. We presented gratings across eight orientations (N = 20 repetitions per orientation) in random order and computed trial-averaged ΔF/F for different orientations. We found strong and widespread visually-evoked activity across the FOV (Fig. 2A-B). To assess visual responsiveness of cells, we calculated single-cell ΔF/F activity traces across visual presentations (Fig. 2C displays 20 example cells across three consecutive trials from one animal). We found many cells respond to drifting grating stimuli (931/1046 cells responsive across N = 3 animals, 2-sample t-test, stimulus vs. baseline, all stimulus directions pooled; p<0.001 threshold; test done with Bonferroni correction across neurons within each experiment). Many neurons also exhibit spontaneous activity (Fig. 2C) between grating presentations.

**Figure 2:**
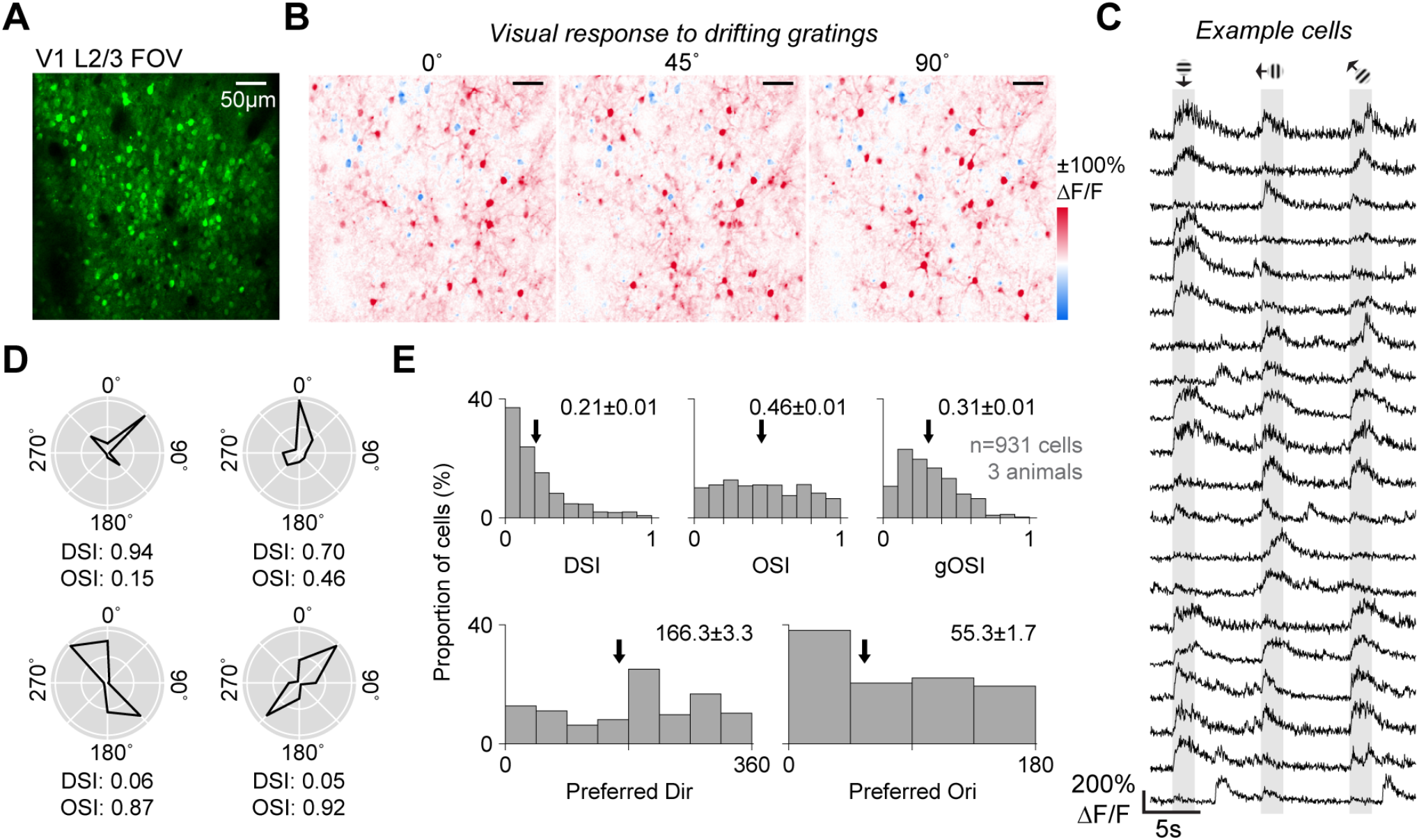
V1 cells expressing jGCaMP8s-P2A-stChrimsonR show expected visually-evoked and spontaneous activity. ***A*** Two-photon calcium imaging FOV (414 × 414 μm) in layer 2/3 of mouse V1. ***B*** Example trial-average visual response maps to retinotopically-aligned 15 degree FWHM Gabor drifting gratings showing evoked activity (ΔF/F) to 0°, 45°, and 90° orientations across the FOV (8 orientations presented in total in random order and lasting for 2 seconds, N = 20 repetitions/orientation). ***C*** ΔF/F traces in example cells (N = 20) across three consecutive trials showing stimulus-evoked responses as well as spontaneous activity between visual presentations. ***D*** Trial-average responses across 8 orientations in four example cells showing direction and orientation tuning. DSI and OSI were calculated using the average activity (ΔF/F) across the 2 second visual period. Polar plots depict responses normalized to peak response across all 8 orientations for each cell. ***E*** Histograms showing distributions of tuning metrics across all cells (N = 931 cells across 3 animals). Black arrows indicate the mean value of each metric. Mean ± SEM of each metric across all animals shown in top right of each histogram. (DSI: direction selectivity index, OSI: orientation selectivity index, gOSI: global orientation selectivity index). This figure: N = 3 mice. **A**-**D**: mouse 3, **E**: mouse 3, 5, and 6.

Next, we examined tuning for the direction and orientation of the stimuli and found the distribution of neurons’ tuning was very similar to what has been previously reported with GCaMP6 or GCaMP7 by other laboratories. For every cell, we calculated selectivity indices for direction (DSI) and orientation (OSI and global OSI, or gOSI) of the drifting grating, as well as preferred direction and orientation (Fig. 2D-E). We found the distributions of these tuning metrics match closely with prior reports using calcium indicators (GCaMP6s, Kondo and Ohki, 2016, see Fig. 3D, Fig. 4D, supp Fig. 7D; GCaMP7s, Bounds et al., 2022, see Fig. 3D). We find that preferred orientations and directions are evenly distributed (Fig. 2E) and tuning index histograms are similar to what has been previously reported. In these reports, mean DSI was 0.27 (Kondo and Ohki, 2016), while our mean DSI was 0.21 ± 0.21 (1 std. dev.) Similarly, their mean OSI was 0.62 (Kondo and Ohki, 2016) and 0.56 (Ai203; Bounds et al., 2022), while we report a mean OSI of 0.46 ± 0.28. For gOSI, they report a mean of 0.46 (Kondo and Ohki, 2016), while we report a mean gOSI of 0.31 ± 0.18. These data demonstrate that expression of jGCaMP8s-P2A-stChrimsonR allows for reliable two-photon measurements of sensory-evoked and ongoing calcium activity.

**Figure 3:**
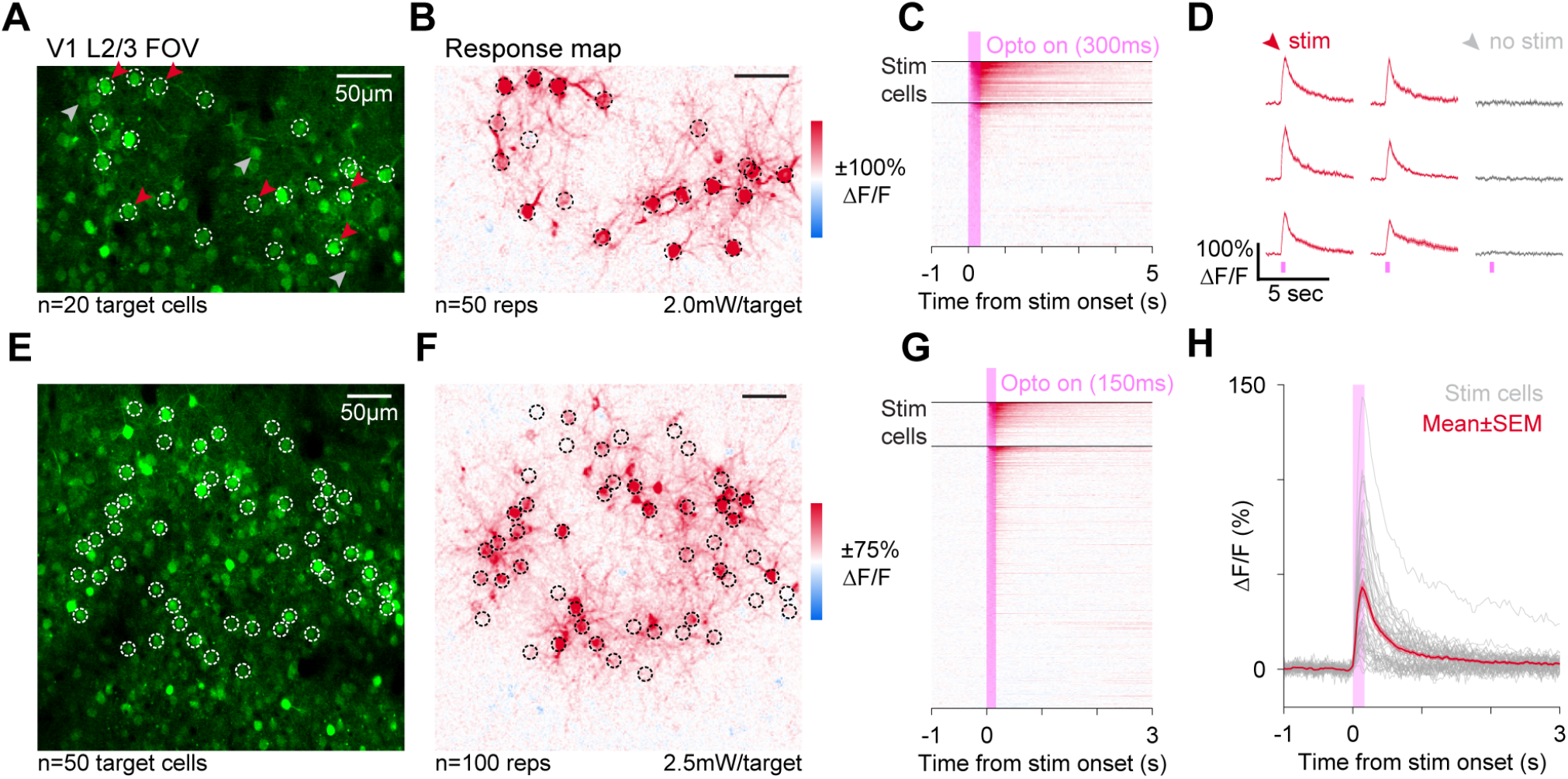
Simultaneous holographic stimulation and calcium imaging in cells expressing jGCaMP8s-P2A-stChrimsonR. ***A*** Two-photon imaging FOV (340 × 210 μm) in mouse V1 at a depth of 122 μm from the pia. Holographic stimulation pattern is denoted by white rings (N = 20 targets). Red and grey arrows represent examples depicted in **D** for stimulated and nearby, non-stimulated cells, respectively. ***B*** Response map in the same FOV as **A** depicting the mean optogenetically-evoked activity (ΔF/F) over multiple stimulations (N = 50 stim reps). Stimulation power was 2.0 mW/target and cells were stimulated for 300 ms each repetition (stimulation delivered during imaging horizontal flyback, reported power is adjusted for this duty cycle, see Methods) using 10 μm diameter disk patterns. ***C*** Trial-averaged activity traces for all cells identified in the FOV (N = 89 total cells). Black horizontal bars delineate stimulation targets (top) from other cells in the FOV (bottom). ***D*** Trial-averaged cell traces (mean ± SEM, n=50 stim reps) for 9 example cells (N = 6 stimulated, red and N = 3 non-stimulated, gray). Errorbars are ±SEM and are small enough to show only as slightly thickened lines. Example traces correspond to cells indicated in **A** by arrows. ***E-G*** Same as **A-C**, in a different FOV in the same animal 1 month later (FOV 414 × 414 μm, depth = 130 μm, N = 50 targets, N = 348 total cells, N = 100 stim reps, stimulation power was 2.5 mW/target for 150 ms using 10 μm diameter disk patterns). ***H*** Trial-average cell traces (N = 100 stim reps) for the 50 targeted cells in **E** (gray lines) and the mean±SEM cell trace (red line). This figure: N = 1 mouse. **A**-**H** mouse 1.

### Holographic pattern stimulation in vivo of cells expressing jGCaMP8s and stChrimsonR

One goal of achieving co-expression of both an indicator and an opsin is to facilitate all-optical interrogation of brain circuits. Therefore, next, to test the ability of cells expressing the construct to respond to optogenetic stimulation, we measured jGCaMP8s activity via two-photon imaging *in vivo* (920 nm imaging wavelength, 15–20 mW imaging power, 80 MHz pulse rate, 0.19–0.25 nJ/pulse) while stimulating cells with holographic light patterns (1030 nm, 500 kHz pulse rate, 13–16 nJ/pulse). We used holography to simultaneously stimulate cells in mouse V1 with 10 μm-diameter light spots in a single depth plane. This stimulation (20 targets, 2.0 mW/target for 300 ms, see Methods for details, Fig. 3A) led to clear optogenetically-evoked responses in targeted cells (Fig. 3B-D). Reliable responses were measured in 19/20 targeted cells (p<0.001; 2-sample t-test with Bonferroni correction for multiple comparisons, between stim and baseline ΔF/F, Methods). Nearby unstimulated cells in the FOV showed minimal photoactivation (Fig. 3C-D). We observed similar results for photostimulation experiments in other mice (30/32 cells across N = 3 mice show reliable responses; p<0.001 threshold, 2-sample t-test with Bonferroni correction for each experiment; Extended Fig. 3-1).

This viral approach allows stimulation of many cells simultaneously (50 targets, 2.5 mW/target for 150 ms, Fig. 3E-H). We found 45/50 targeted cells showed significant activation (although some cells are activated more strongly than others, Fig. 3H and Extended Fig. 3-1, Methods for response reliability measure). However, when using 50 stimulation targets, we find more off-target activation in nearby cells (Fig. 3G), when compared to 20 stimulation targets (Fig. 3C). It seems possible that this increase in off-target activation is due to network effects, such as postsynaptic summation, but we did not explore this further, as our primary goal was to determine whether a large proportion of targeted neurons could be activated reliably.

We find that a majority of jGCaMP8s-expressing cells selected for holographic targeting show consistent responses to repeated photostimulation, indicating these cells are successfully co-expressing both indicator and opsin.

## Discussion

In order to facilitate two-photon holographic photostimulation experiments with simultaneous calcium imaging, we designed a Cre-based, single viral approach to express both opsin and indicator in Cre-expressing cells of interest. The single viral approach simplifies experimental preparation by eliminating factors related to relative concentrations of opsin and indicator. It also effectively addresses challenges with co-expression of opsin and indicator in the same cells that can occur when using multiple viruses or combinations of viruses and transgenic animals. Our results indicate that cells expressing AAV9-hSyn-DIO-jGCaMP8s-P2A-stChrimsonR exhibit strong and consistent responses to targeted holographic photostimulation. Also, visual responses measured using this approach have population distributions similar to those previously reported with expression of calcium sensors, indicating normal visual function is maintained and neural activity can be reliably recorded.

### Robust activation of neurons with ChrimsonR likely reflects changes in firing rates, instead of precisely timed single spikes

A central appeal of two-photon holography is the ability to dynamically identify and select groups of cells for photostimulation based on functional properties. However, a limiting factor for such experiments is achieving reliable co-expression of opsin and indicator in many cells throughout the tissue. Addressing this limitation has been a core focus of past genetic approaches: successful co-expression yields the ability to measure responses to optogenetic stimulation. Recent studies typically find many, but not all, cells are responsive to photostimulation (for example, ∼80% with Ai203 transgenic line, Bounds et al., 2022; ∼90% with ChRmine, ∼85% with ChRome2s, ∼75% ChRome2f, Sridharan et al., 2022). On par with prior reports for other opsins, we found similar and greater levels of photoactivatable cells in our experiments (∼95% of cells photoactivatable). However, our data differ from previous reports with ChrimsonR, which suggest very few ChrimsonR cells are reliably activated with two-photon stimulation (∼10%, see Fig. 7C-D of Sridharan et al., 2022). However, this report aimed to induce precisely timed single spikes, and thus used much stronger stimulation intensities (∼13x greater than in our experiments, their work 0.41 mW/μm^2^, here 0.03 mW/μm^2^) and much shorter stimulation pulse lengths (5 ms, repeated 5 times at 30 Hz). Given the relatively smaller photocurrents produced by ChrimsonR activation as compared to other opsins (see Fig 1G of Sridharan et al., 2022), it might be unsurprising they were unable to activate many neurons with these small pulses.

In fact, it might be argued that smaller stimulation currents, which modulate the firing rate of neurons without reliably driving single spikes with precise timing, are similar to inputs neurons receive during many forms of visual sensory stimulation. During flashed or drifting grating stimuli (Niell and Stryker, 2008; Busse et al., 2011; Glickfeld et al., 2013), mouse neurons change their firing rates by only up to tens of spikes per second or less, and fire with irregular, Poisson-like timing. Electrophysiological experiments show that activation of stChrimsonR in the visual cortex with longer pulses produces an elevation of neurons’ firing rates while neurons’ firing remains irregular, as occurs with sensory stimuli (O’Rawe et al., 2022). The elevated firing rate continues for as long as cells are illuminated (up to 1000 ms).

In sum, the difference between past low fractions of activated cells with ChrimsonR and the large fractions of excitable cells we show here may be due to a difference in stimulation duration and thus total current injected. Longer pulses should be expected to produce more spikes on average and improve the chances of detecting photoactivation, enabling flexible targeting of behaviorally-relevant neurons. Moreover, by pairing ChrimsonR with jGCaMP8s, an indicator optimized for the detection of single spikes (with the tradeoff of becoming nonlinear with fewer spikes than jGCaMP8m or 8f; see Supp Fig. 6 from Zhang et al., 2021), we improve our ability to detect photoactivation, while prior reports have paired ChrimsonR with the less-sensitive GCaMP6 variant.

### Cell responses to both optogenetic stimulation and visual stimulation are robust and stable over time

Many or all neurons transfected with our virus show GCaMP fluorescence throughout the cell instead of being excluded from the nucleus. In single-promoter GCaMP expression, cell filling with GCaMP is often associated with a pathological state which develops over time and leads to bright fluorescence no longer modulated by activity (Tian et al., 2009; Chen et al., 2013; Packer et al. 2015). In this study, however, most or all neurons are filled, but the filled neurons are neither extremely bright, nor static. In fact, our two-photon stimulation results (Fig. 3) and our visual stimulation results (Fig. 2) suggest that the filled neurons are responding to input and reflecting these changes in GCaMP fluorescence. The filling might arise because the cleavage at the P2A site after transcription may yield a GCaMP protein that is trafficked slightly differently than the GCaMP constructs previously expressed with a single promoter. However, the neural responses we measured with this virus were robust, stable, and were similar to sensory-evoked responses previously measured in other work (Fig. 2). Therefore, if there is some difference in protein trafficking, it seems to leave the GCaMP responses to neural activity intact.

### Future and conclusion

Currently, most studies employing all-optical methods for circuit dissection have focused on cortical areas. However, holographic stimulation can be used to investigate network function in other brain regions, as in the olfactory bulb (Gill et al., 2020) or the basolateral amygdala (Piantadosi et al., 2022). While viral tropisms may affect whether any given virus that works well in the cortex also works well in other brain regions, bicistronic delivery methods of both opsin and indicator will simplify testing of expression strategies.

In the present work, we build on previous efforts using separate viruses to express various GCaMP indicators (GCaMP6f, GCaMP6s, jGCaMP7s) with ChrimsonR (in mice, Attinger et al., 2017; Stamatakis et al., 2018; Chettih and Harvey, 2019; Gill et al., 2020; Daie et al., 2021; Akitake et al., 2022; O’Rawe et al., 2022; and in zebrafish, Förster et al., 2017). Our bicistronic virus adds to the growing genetic toolbox enabling future experiments requiring flexible patterned stimulation methods by offering a single viral approach to express the latest GCaMP8s indicator alongside the ChrimsonR opsin in cells of interest.

## Materials and methods

### Virus

Both pAAV-hSyn-DIO-ChrimsonR-mRuby2-ST (Addgene Plasmid #105448, RRID:Addgene 105448) and pGP-AAV-syn-jGCaMP8s-WPRE (Addgene Plasmid #162374, RRID:Addgene_162374) plasmids were used to build the pAAV-hSyn-DIO-jGCaMP8s-P2A-stChrimsonR construct. The mRuby fluorescent tag from pAAV-hSyn-DIO-ChrimsonR-mRuby2-ST was removed, and the sequence encoding jGCaMP8s was cloned into the construct along with a P2A peptide linker. The plasmid was used for packaging into an adeno-associated virus (AAV9). The plasmid version of this construct will be available on Addgene (Addgene Plasmid #174007, RRID:Addgene 174007).

### Animals and Surgery

All animal procedures were performed in accordance with the NIH Institutional Animal Care and Use Committee’s (IACUC) regulations. *Emx1-Cre* mice (The Jackson Laboratory; RRID:IMSR JAX:005628) were used in all experiments to target expression of Cre to excitatory, glutamatergic neurons. N = 6 total animals were used in this study (N = 3 male, N = 3 female); no differences due to sex were noted in the results. Mice 2 months of age or older were anesthetized with isoflurane (1–3% in 100% O_2_ at 1 L/min) and kept on a heating pad for warmth. An intraperitoneal injection of dexamethasone (3.2 mg/kg) was administered before incision to reduce inflammation. The skull was exposed, and a custom metal head post was positioned at the base of the skull. A 3mm diameter circular craniotomy was made over the left hemisphere of primary visual cortex (ML -3.1 mm, AP +1.5 mm relative to Lambda) using an air driven dental drill (Aseptico; Woodinville, WA) with a Neoburr drill bit (Friction Grip 1/4; Microcopy; Kennesaw, GA). AAV9-hSyn-DIO-jGCaMP8s-P2A-stChrimsonR was diluted in phosphate-buffered saline to either a final titer of 2.62×10^12^ GC/mL (mouse 1, 3, 5, and 6), 3.37×10^12^ (mouse 2), or 4.72×10^12^ (mouse 4), and 2 nmol sulforhodamine 101 was added to visualize injection progression in the brain. Virus was injected unilaterally with a stereotactic syringe pump (Stoelting; Wood Dale, IL) through a pulled glass pipette tip cut to an opening of 10–15 μm diameter. Injections were targeted to 200 μm below the surface of the brain and administered at a rate of 0.1 μL/min for a total volume of 300 nL per injection site (5–10 injection sites). A 3 mm optical window (Tower Optical; Boynton Beach, FL) was implanted over the craniotomy. Both the optical window and metal head post were fixed to the skull using C&B metabond dental cement dyed black (Parkell; Edgewood, NY). Last, a custom-made removable light-blocking cover was fixed atop the implant to prevent ambient light exposure to the opsin. Animals were individually housed after surgery, and because the present study does not include behavioral assays, individual housing is unlikely to significantly impact our results. Mice were imaged three or more weeks post-injection. All animals were housed in a 12:12 hour reverse light-dark cycle and allowed food and water ad libitum.

### Widefield fluorescence imaging

Widefield fluorescence imaging was done using a Discovery stereo microscope (Zeiss; Jena, Germany) with an X-Cite XYLIS LED source (Excelitas; Mississauga, Canada) and a blue excitation and green emission filter set (KSC XXX-814; Kramer Scientific; Amesbury, MA). Images were collected (200 ms exposure period) using a Retiga R3 CCD camera (QImaging, Inc.; Surrey, BC).

### *In vivo* two-photon calcium imaging

To perform two-photon calcium imaging, animals were first head-fixed under a 16x water-immersion objective (Nikon; Tokyo, Japan). Imaging was performed using a custom-built microscope using MIMMS (Modular In vivo Multiphoton Microscopy System) components (Sutter Instruments; Novato, CA) and a Chameleon Discovery NX tunable femtosecond laser (Coherent, Inc.; Santa Clara, CA). Imaging was controlled using ScanImage software (MBF Biosciences; Williston, VT) in MATLAB. A small volume (∼1 mL) of clear ultrasound gel was placed over the optical window to immerse the objective. Calcium responses were measured at approximately 100–200 μm below the surface of the pia in layers 2/3 of primary visual cortex using a 410 × 410 μm field of view, except where noted.Imaging was performed via bidirectional raster scanning with a resonant-galvo system (8 kHz resonant scanner, 512 lines, ∼30 Hz frame rate) using 920 nm wavelength light at 15–20 mW measured at the front aperture of the objective (pulse rate 80 MHz, pulse energy 0.19–0.25 nJ/pulse).

### *In vivo* two-photon holographic photostimulation

Holographic photostimulation was performed using a Satsuma femtosecond pulsed laser (Amplitude Laser; Pessac, France) at 1030 nm wavelength along a second optical path (galvo-galvo) integrated into the two-photon microscope just before the tube lens using a polarizing beam combiner. A spatial light modulator, or SLM (1920 × 1152 pixels; Meadowlark Optics, Frederick, CO), was used to generate holographic patterns of 10 μm diameter disks within a 2-dimensional plane (aligned to the imaging focal plane). The SLM was followed by a relay lens system (two achromatic lenses, with focal lengths 250 mm and 100 mm) in a 4-f configuration between SLM and galvanometers. The zero order (undiffracted beam) from the SLM was blocked using a small amount of furnace cement (30–40 μm diameter) on a glass slide. Stimulation targets were defined and SLM phase masks were computed using ScanImage software (MBF Biosciences; Williston, VT). Stimulation was applied for either 150 ms at 6.5 mW/target (Fig. 3A-D) or 300 ms at 8mW/target (Fig. 3E-H) with a 500 kHz pulse rate (pulse energy 13 or 16 nJ/pulse). The laser was gated on during horizontal flyback periods and off during the imaging pixel acquisition (on time 19 μs, off time 44 μs, duty cycle 30%) to allow for imaging of responses during stimulation periods (imaging frame clock was inverted via a TTL logic gate; Pulse Research Lab; Torrance, CA.) Stimulation power was thus reduced by a factor of 0.3 from laser power measured without this gating, resulting in effective stimulation powers of 2.0 mW/target (Fig. 3A-D) and 2.5 mW/target (Fig. 3E-H).

### Retinotopic Mapping

To map the retinotopic position of visual stimuli in V1 under the optical window, we performed hemodynamic intrinsic imaging in awake head-fixed animals. We presented visual stimuli (drifting square wave gratings, 0.1 cycles/degree, 10 degrees diameter) for 5 seconds (with 10 seconds between presentations) at different retinotopic positions and measured reflected 530 nm light on the brain to quantify hemodynamic-related changes in absorption. The 530 nm light was delivered using a fiber-coupled LED (M530F2; Thorlabs, Newton, NJ) and imaging was collected on the same stereo microscope used for widefield fluorescence imaging using a 1x widefield objective and a green long-pass emission filter. Imaging was acquired at 2 Hz. Changes in reflectance were computed for every stimulus location between a baseline period (5 seconds prior to stimulus onset) and a response period (2.5 second window starting 3 seconds after stimulus onset). A centroid of the hemodynamic response was computed for each stimulus location and an average retinotopic map was fit to the positions of the centroids. Retinotopic maps were then used to guide stimulus locations for two-photon imaging measures of visual responses to Gabor stimuli.

### Visual stimulation

To measure visual responses during two-photon imaging, we presented awake animals with Gabor patches (sinusoidal drifting gratings filtered with a gaussian mask with 15 degree full-width half-max) with spatial frequency 0.1 cycles/degree for 2 second periods (6 seconds of gray screen between presentations) at 100% contrast. An LCD monitor with neutral gray background was used to present visual stimuli and was positioned approximately 20 centimeters in front of the animal. Drifting gratings of 8 different directions (45 degree increments) were presented in random order across trials. Each direction was presented for 20 repetitions.

### Two-photon calcium imaging analysis

Two-photon calcium imaging data was first downsampled from 512 × 512 pixels to 256 × 256 to ease handling of data. Background correction was performed by computing the average intensity image across frames and subtracting the minimum pixel value of this average from the image stack. All remaining negative pixel values (due to noise) were then set to zero. We motion corrected all images using the CaImAn toolbox (Giovannucci et al., 2019) and performed cell segmentation using Suite2p to allow manual selection of cell masks (Pachitariu et al., 2017). Fluorescence intensity traces were then calculated as the average intensity across all pixels within a cell’s segmented mask. To quantify cell activity, we computed ΔF/F_0_ for each cell. F_0_ was defined as the average fluorescence across the 50 imaging frames that occurred directly prior to stimulus presentation for all trials (N = 160 trials, Fig. 2; N = 50 trials, Fig. 3C; N = 100 trials, Fig. 3G) in an imaging session.

Reliability measures for photostimulation responses were quantified by performing a 2-sample t-test (with Bonferroni correction for multiple comparisons within each experiment, p<0.001) using ΔF/F_0_ values between the photostimulation period (N = 5 frames, Fig. 3B; N = 10 frames, Fig. 3F) and an equivalent number of frames preceding stimulus onset across all trials. Visually-responsivity was quantified in the same fashion using the full visual stimulus period (N = 60 frames).

For visual display of ΔF/F responses across the FOV (as in Fig. 2B, Fig. 3B, and Extended Fig. 3-1B), F_0_ was computed for every pixel in the same manner as for cell-based calculations. We then smoothed the F_0_ image using a gaussian filter (**σ** = 20, radius = 80.5 pixels) to act as a means of local contrast adaptation. F was computed at every pixel as the average value across a response window (N = 60 frames, Fig. 2B; N = 10 frames, Fig. 3B and Extended Fig. 3-1B; N = 15 frames, Fig. 3F) following stimulation onset and across all trials (for Fig. 2B, N = 20 trials/stimulus direction).

### Visual tuning analysis

To quantify visual tuning in individual cells, we computed a direction selectivity index (DSI), orientation selectivity index (OSI), and global orientation selectivity index (gOSI) for all visually-responsive cells following the methods of Kondo and Ohki, 2016. For each metric, 0 indicated no selectivity and 1 indicated maximal selectivity. We first calculated a tuning curve for each cell as the average ΔF/F_0_ activity across the entire visual stimulus period (N = 60 frames, or 2 seconds) and across all trials for each of the 8 stimulus directions (N = 20 trials/direction). The stimulus direction corresponding to the peak value of this tuning curve was determined to be the preferred direction for a cell. The response at this direction, R_prefDir_, as well as the opposite direction (180 degrees away), R_oppoDir_, was used to calculate the DSI as:

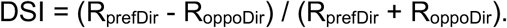

To calculate OSI, we first averaged opposite pairs of directions of the tuning curve to yield average responses to each of the 4 stimulus orientations. The preferred orientation of cells was determined to be the peak value between the 4 orientations. The response at the preferred orientation, R_prefOri_, as well as the response at the orthogonal orientation (90 degrees away), R_orthOri_, was used to calculate the OSI as:

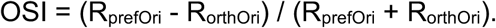

Last, the gOSI for each cell was computed via a vector averaging method (Swindale, 1998):

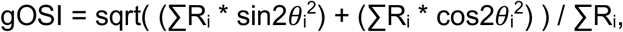

where *i* represents the *i*th direction (equivalent to 1 - circular variance).

## Author contributions

PKL, BA, and MH designed the construct. ZZ performed viral injections and surgical implants. PKL collected two-photon data. ZZ collected widefield fluorescence data. PKL analyzed the data. YD and MH designed and constructed the two-photon microscope and two-photon holographic stimulation equipment. PKL, NGF, and MH wrote the manuscript.

## Acknowledgments

We thank Georg Jaindl and others from Vidrio/MBF Biosciences for expert assistance and support, Victoria M. Scott for assistance in virus testing, and Tommaso Fellin for comments on the manuscript. This work was supported by the NIH BRAIN Initiative, grant U19NS107464, and the NIMH Intramural Research Program, ZIAMH0020967.

## Extended Figures

**Extended Figure 1-1:**
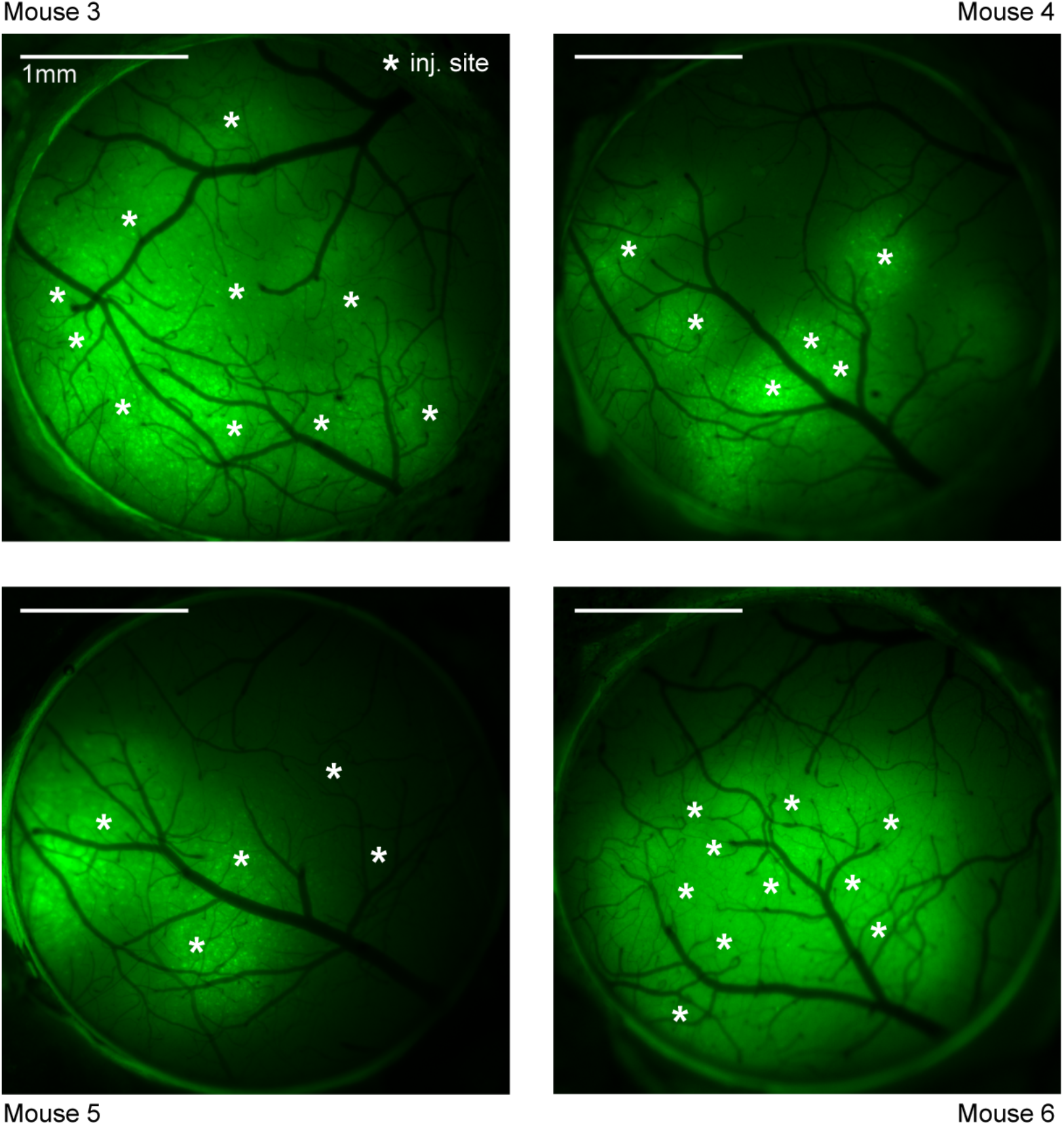
Widefield fluorescence imaging of jGCaMP8s-P2A-stChrimsonR expression. Widefield fluorescence imaging through 3 mm optical windows in experimental animals. White stars denote injection sites for each animal (Mouse 3: N = 10 sites, mouse 4: N = 6 sites, mouse 5: N = 5 sites, mouse 6: n = 10 sites). Images were acquired 22-53 days after injection.

**Extended Figure 3-1:**
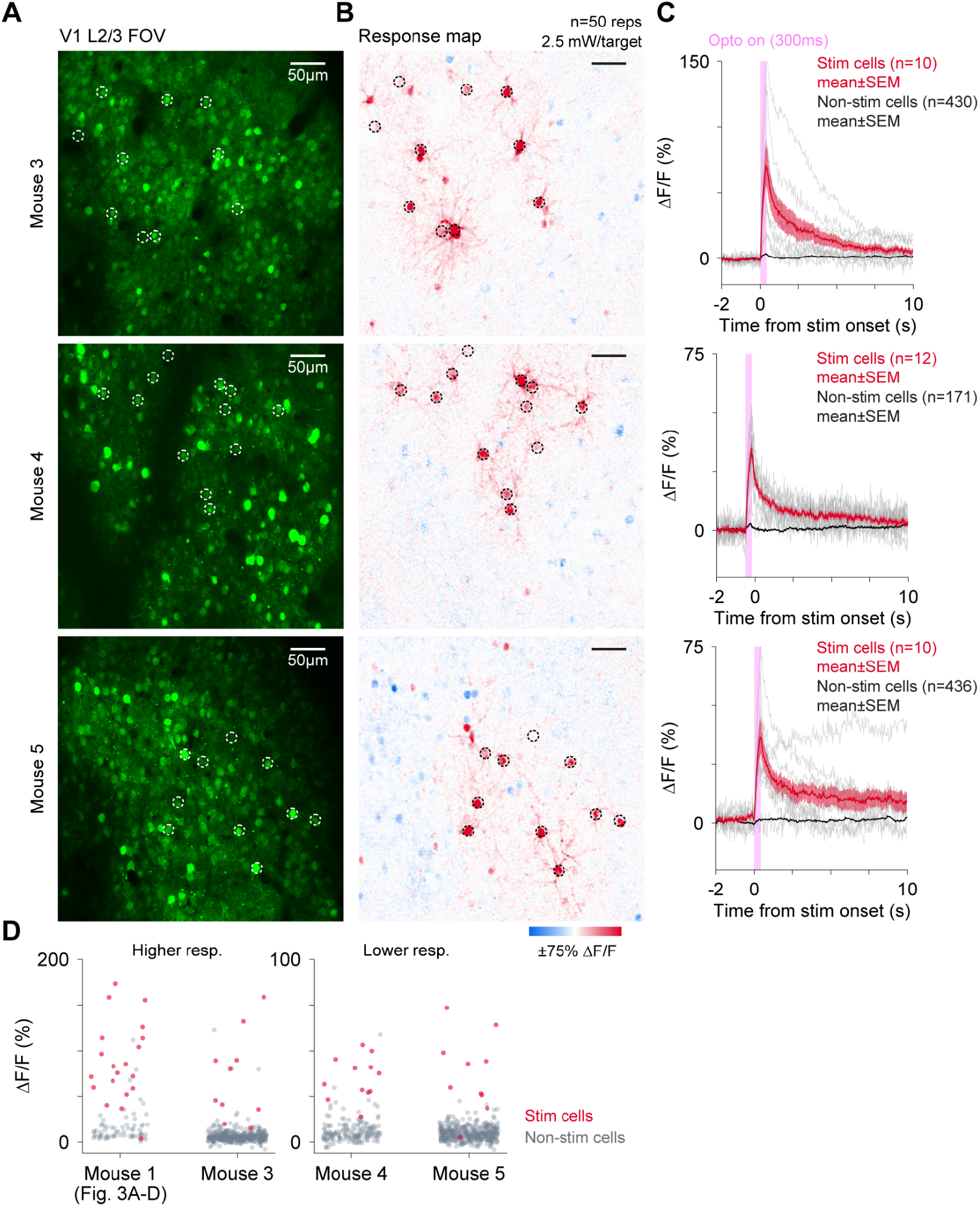
Holographic stimulation of cells expressing jGCaMP8s-P2A-stChrimsonR. ***A*** Two-photon imaging FOV (410 × 410 μm) in mouse V1 in 3 example mice at a depth of 130 μm (top), 140 μm (middle), and 110 μm (bottom). Holographic stimulation pattern is denoted by white rings (top: N = 10 targets, middle: N = 12 targets, bottom: N = 10 targets). ***B*** Corresponding stimulation response maps showing mean optogenetically-evoked activity (ΔF/F) over multiple stimulation repetitions (top: N = 50 repetitions, middle: N = 50, bottom: N = 58). Stimulation was applied at 2.5 mW/target for 300 ms on each repetition using 10 μm diameter disk patterns. ***C*** Trial-averaged activity of all stimulated cells (light gray lines) plotted with mean ± SEM activity across all stimulated cells (red lines) and all non-stimulated cells (black lines). SEM is present for non-stimulated cells, but small relative to the size of the plotted line. Top: 10/10 responsive cells, middle: 12/12 responsive cells, bottom: 9/10 responsive cells (p<0.001; 2-sample t-test with Bonferroni correction (see Methods for details). ***D*** Trial-averaged peak responses to photostimulation (from last 150 ms of stimulation) in all cells for each experiment (including from Fig. 3A-D). Red points: cells targeted for stimulation, gray points: non-stimulated cells. Note different y-axes for mouse 1, 3 (higher responses) versus mouse 4, 5 (lower responses). Some possible explanations for variations in magnitude of photostimulation responses between animals may be differences in injection titers (mouse 1, 3, and 5: 2.6×10^12^ GC/mL; mouse 4: 4.7×10^12^ GC/mL) and/or optical window quality.

## Notes

### Competing Interest Statement

The authors have declared no competing interest.

## References

Adesnik H, Abdeladim L (2021) Probing neural codes with two-photon holographic optogenetics. Nat Neurosci 24:1356–1366.

Akitake B, Douglas HM, LaFosse PK, Deveau CE, Li AJ, Ryan LN, Duffy SP, Zhou Z, Deng Y, Histed MH (2022) Amplified cortical neural responses as animals learn to use novel activity patterns. bioRxiv:2022.07.10.499496.

Attinger A, Wang B, Keller GB (2017) Visuomotor Coupling Shapes the Functional Development of Mouse Visual Cortex. Cell 169:1291–1302.e14.

Bounds HA, Sadahiro M, Hendricks WD, Gajowa M, Gopakumar K, Quintana D, Tasic B, Daigle TL, Zeng H, Oldenburg IA, Adesnik H (2022) Ultra-precise all-optical manipulation of neural circuits with multifunctional Cre-dependent transgenic mice. bioRxiv:2021.10.05.463223.

Busse L, Ayaz A, Dhruv NT, Katzner S, Saleem AB, Scholvinck ML, Zaharia AD, Carandini M (2011) The Detection of Visual Contrast in the Behaving Mouse. Journal of Neuroscience 31:11351– 11361.

Cardin JA, Carlén M, Meletis K, Knoblich U, Zhang F, Deisseroth K, Tsai L-H, Moore CI (2009) Driving fast-spiking cells induces gamma rhythm and controls sensory responses. Nature 459:663–667.

Carrillo-Reid L, Han S, Yang W, Akrouh A, Yuste R (2018) Triggering visually-guided behavior by holographic activation of pattern completion neurons in cortical ensembles. bioRxiv:394999.

Carrillo-Reid L, Han S, Yang W, Akrouh A, Yuste R (2019) Controlling Visually Guided Behavior by Holographic Recalling of Cortical Ensembles. Cell 178:447–457.e5.

Chen I-W, Ronzitti E, Lee BR, Daigle TL, Dalkara D, Zeng H, Emiliani V, Papagiakoumou E (2019) In Vivo Submillisecond Two-Photon Optogenetics with Temporally Focused Patterned Light. J Neurosci 39:3484–3497.

Chen T-W, Wardill TJ, Sun Y, Pulver SR, Renninger SL, Baohan A, Schreiter ER, Kerr RA, Orger MB, Jayaraman V, Looger LL, Svoboda K, Kim DS (2013) Ultrasensitive fluorescent proteins for imaging neuronal activity. Nature 499:295–300.

Chettih SN, Harvey CD (2019) Single-neuron perturbations reveal feature-specific competition in V1. Nature 567:334–340.

Daie K, Svoboda K, Druckmann S (2021) Targeted photostimulation uncovers circuit motifs supporting short-term memory. Nat Neurosci 24:259–265.

Daigle TL et al. (2018) A Suite of Transgenic Driver and Reporter Mouse Lines with Enhanced Brain-Cell-Type Targeting and Functionality. Cell 174:465–480.e22.

Dalgleish HW, Russell LE, Packer AM, Roth A, Gauld OM, Greenstreet F, Thompson EJ, Häusser M (2020) How many neurons are sufficient for perception of cortical activity? Elife 9.

Denk W, Delaney KR, Gelperin A, Kleinfeld D, Strowbridge BW, Tank DW, Yuste R (1994) Anatomical and functional imaging of neurons using 2-photon laser scanning microscopy. J Neurosci Methods 54:151–162.

Emiliani V, Cohen AE, Deisseroth K, Häusser M (2015) All-Optical Interrogation of Neural Circuits. J Neurosci 35:13917–13926.

Forli A, Pisoni M, Printz Y, Yizhar O, Fellin T (2021) Optogenetic strategies for high-efficiency all-optical interrogation using blue-light-sensitive opsins. Elife 10.

Förster D, Maschio MD, Laurell E, Baier H (2017) An optogenetic toolbox for unbiased discovery of functionally connected cells in neural circuits. Nature Communications 8.

Gill JV, Lerman GM, Zhao H, Stetler BJ, Rinberg D, Shoham S (2020) Precise Holographic Manipulation of Olfactory Circuits Reveals Coding Features Determining Perceptual Detection. Neuron 108:382–393.e5.

Giovannucci A, Friedrich J, Gunn P, Kalfon J, Brown BL, Koay SA, Taxidis J, Najafi F, Gauthier JL, Zhou P, Khakh BS, Tank DW, Chklovskii DB, Pnevmatikakis EA (2019) CaImAn an open source tool for scalable calcium imaging data analysis. eLife 8.

Glickfeld LL, Histed MH, Maunsell JHR (2013) Mouse primary visual cortex is used to detect both orientation and contrast changes. J Neurosci 33:19416–19422.

Kim JH, Lee S-R, Li L-H, Park H-J, Park J-H, Lee KY, Kim M-K, Shin BA, Choi S-Y (2011) High cleavage efficiency of a 2A peptide derived from porcine teschovirus-1 in human cell lines, zebrafish and mice. PLoS One 6:e18556.

Klapoetke NC et al. (2014) Independent optical excitation of distinct neural populations. Nat Methods 11:338–346.

Kondo S, Ohki K (2016) Laminar differences in the orientation selectivity of geniculate afferents in mouse primary visual cortex. Nat Neurosci 19:316–319.

Mardinly AR, Oldenburg IA, Pégard NC, Sridharan S, Lyall EH, Chesnov K, Brohawn SG, Waller L, Adesnik H (2018) Precise multimodal optical control of neural ensemble activity. Nat Neurosci 21:881–893.

Marshel JH, Kim YS, Machado TA, Quirin S, Benson B, Kadmon J, Raja C, Chibukhchyan A, Ramakrishnan C, Inoue M, Shane JC, McKnight DJ, Yoshizawa S, Kato HE, Ganguli S, Deisseroth K (2019) Cortical layer–specific critical dynamics triggering perception. Science 365:eaaw5202.

Niell CM, Stryker MP (2008) Highly selective receptive fields in mouse visual cortex. J Neurosci 28:7520–7536.

O’Rawe JF, Zhou Z, Li AJ, LaFosse PK, Goldbach HC, Histed MH (2022) Excitation creates a distributed pattern of cortical suppression due to varied recurrent input. bioRxiv:2022.08.31.505844.

Pachitariu M, Stringer C, Dipoppa M, Schröder S, Federico Rossi L, Dalgleish H, Carandini M, Harris KD (2017) Suite2p: beyond 10,000 neurons with standard two-photon microscopy. bioRxiv:061507.

Packer AM, Peterka DS, Hirtz JJ, Prakash R, Deisseroth K, Yuste R (2012) Two-photon optogenetics of dendritic spines and neural circuits. Nat Methods 9:1202–1205.

Packer AM, Russell LE, Dalgleish HWP, Häusser M (2015) Simultaneous all-optical manipulation and recording of neural circuit activity with cellular resolution in vivo. Nat Methods 12:140–146.

Pégard NC, Mardinly AR, Oldenburg IA, Sridharan S, Waller L, Adesnik H (2017) Three-dimensional scanless holographic optogenetics with temporal focusing (3D-SHOT). Nat Commun 8:1228.

Piantadosi SC, Zhou ZC, Pizzano C, Pedersen CE, Nguyen TK, Thai S, Stuber GD, Bruchas MR (2022) Holographic stimulation of opposing amygdala ensembles bidirectionally modulates valence-specific behavior. bioRxiv:2022.07.11.499499.

Prakash R, Yizhar O, Grewe B, Ramakrishnan C, Wang N, Goshen I, Packer AM, Peterka DS, Yuste R, Schnitzer MJ, Deisseroth K (2012) Two-photon optogenetic toolbox for fast inhibition, excitation and bistable modulation. Nat Methods 9:1171–1179.

Rickgauer JP, Tank DW (2009) Two-photon excitation of channelrhodopsin-2 at saturation. Proc Natl Acad Sci U S A 106:15025–15030.

Robinson NTM, Descamps LAL, Russell LE, Buchholz MO, Bicknell BA, Antonov GK, Lau JYN, Nutbrown R, Schmidt-Hieber C, Häusser M (2020) Targeted Activation of Hippocampal Place Cells Drives Memory-Guided Spatial Behavior. Cell 183:2041–2042.

Rowland JM, van der Plas TL, Loidolt M, Lees RM, Keeling J, Dehning J, Akam T, Priesemann V, Packer AM (2021) Perception and propagation of activity through the cortical hierarchy is determined by neural variability. bioRxiv:2021.12.28.474343.

Russell LE, Dalgleish HWP, Nutbrown R, Gauld OM, Herrmann D, Fişek M, Packer AM, Häusser M (2022) All-optical interrogation of neural circuits in behaving mice. Nat Protoc 17:1579–1620.

Schnütgen F, Doerflinger N, Calléja C, Wendling O, Chambon P, Ghyselinck NB (2003) A directional strategy for monitoring Cre-mediated recombination at the cellular level in the mouse. Nat Biotechnol 21:562–565.

Shemesh OA, Tanese D, Zampini V, Linghu C, Piatkevich K, Ronzitti E, Papagiakoumou E, Boyden ES, Emiliani V (2017) Temporally precise single-cell-resolution optogenetics. Nat Neurosci 20:1796–1806.

Sridharan S, Gajowa MA, Ogando MB, Jagadisan UK, Abdeladim L, Sadahiro M, Bounds HA, Hendricks WD, Turney TS, Tayler I, Gopakumar K, Oldenburg IA, Brohawn SG, Adesnik H (2022) High-performance microbial opsins for spatially and temporally precise perturbations of large neuronal networks. Neuron 110:1139–1155.e6.

Stamatakis AM, Schachter MJ, Gulati S, Zitelli KT, Malanowski S, Tajik A, Fritz C, Trulson M, Otte SL (2018) Simultaneous Optogenetics and Cellular Resolution Calcium Imaging During Active Behavior Using a Miniaturized Microscope. Front Neurosci 12:496.

Swindale NV (1998) Orientation tuning curves: empirical description and estimation of parameters. Biol Cybern 78:45–56.

Szymczak AL, Workman CJ, Wang Y, Vignali KM, Dilioglou S, Vanin EF, Vignali DAA (2004) Correction of multi-gene deficiency in vivo using a single “self-cleaving” 2A peptide–based retroviral vector. Nat Biotechnol 22:589–594.

Tian L, Hires SA, Mao T, Huber D, Chiappe ME, Chalasani SH, Petreanu L, Akerboom J, McKinney SA, Schreiter ER, Bargmann CI, Jayaraman V, Svoboda K, Looger LL (2009) Imaging neural activity in worms, flies and mice with improved GCaMP calcium indicators. Nat Methods 6:875– 881.

Zhang Y et al. (2021) Fast and sensitive GCaMP calcium indicators for imaging neural populations. bioRxiv:2021.11.08.467793.

